# A simple approach to multiplexed PCR amplicon sequencing of plumage-associated loci in *Vermivora* warblers

**DOI:** 10.1101/2025.02.03.636269

**Authors:** David P. L. Toews, Catharine E. Besch

## Abstract

The dynamics of hybridization between golden-winged (*Vermivora chrysoptera*) and blue-winged warblers (*V. cyanoptera*) has been of interest for over a century. Whole genome analysis found only a small number of genomic regions that differed between the species. We previously developed a restriction enzyme-based RFLP approach to genotype large numbers of individuals at each of these loci. Here we extend this approach to an amplicon sequencing method to genotype individuals at these six plumage-associated regions. We demonstrate the efficacy using preliminary data from 4 golden-winged and 4 blue-winged warblers as well as provide the data and scripts necessary to analyze these data for other interested in replicating this approach. Our hope is that these data are useful for other researchers interested in genotyping *Vermivora* warblers.

**Additional methods:** https://github.com/david-toews/vermivora_amplicon

## Introduction

The dynamics of hybridization between golden-winged (*Vermivora chrysoptera*) and blue-winged warblers (*V. cyanoptera*) has been of interest for over a century (Faxon 1913). Our previous worked initially sequencing whole genomes of the two species identified only six regions that were divergent between them within an otherwise homogeneous genomic background. Within nearly all of these regions housed genes involved in different aspects of plumage pigmentation—including melanogenesis and carotenoid-based traits—presumably underlying the phenotypic differences between the taxa (Toews et al. 2016, Baiz et al. 2020, Baiz et al. 2021).

We previously developed a restriction enzyme-based RFLP approach to genotype large numbers of individuals at each of these loci. However, the time involved in running—and sometimes re-running—agarose gels to obtain individual genotypes for hundreds of birds across the six loci was significant. Given the advances and accessibility of genomic analysis and amplicon sequencing, we therefore endeavored to optimize a method to genotype hundreds of individuals at the same time using amplicon sequencing. Given there are a number of research labs interested in the hybridization dynamics in *Vermivora*—and have been using the RFLP method—our goal here is to provide a simple outline of our methods such that others can replicate the same amplicon sequencing approach, if desired.

## Methods

### Sample and library preparation

For a proof of concept, we obtained blood samples from 8 *Vermivora* warblers, 4 phenotypic golden-winged warblers and 4 phenotypic blue-winged warblers. From each bird, we collected blood samples from the brachial vein and stored them in Queen’s lysis buffer. We conducted DNA extractions using the Qiagen DNeasy Blood and Tissue kit following the nucleated blood spin-column protocol. We used six primers (Table S1 from Toews et al. 2016) and prepared a single primer mix containing all forward and reverse primers each at a final concentration of 0.7uM. We ordered the six previously developed primers with P5 [TCGTCGGCAGCGTCAGCTGTGTATAAGAGACAG] and P7 [GTCTCGTGGGCTCGGAGATGTGTATAAGAGACAG] adapter overhangs (note there is an error in Table S1 in Toews et al. [2016] regarding the restriction enzymes cut sites, which should show that locus “24-563” cuts GW – C with BW – A, and locus “25-653” cuts BW – T, with GW – C).

For each sample, we conducted individual 15uL PCR reactions containing: 3 uL 5X Platinum Buffer (Invitrogen 14966005), 0.3 uL 10mM dNTP mix (Promega U151A), 1.5 uL of the pooled primer mix, 0.12 uL of Platunim Taq HS II (Invitrogen 14966005), 8.58 uL of nuclease-free water, and 1.5 uL of the DNA samples (Qubit concentrations: 22.6-62.4 ng/uL). All primers were previously designed to have an annealing temperature of ∼60^°^C.

The thermocycling protocol consisted of:

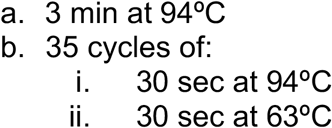

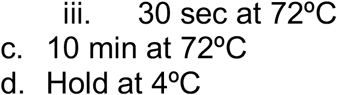

For the indexing PCR, we initially diluted PCR products by 10X. We then added combinatorial indexes based on Faircloth and Glenn (2012). The 30 uL indexing reaction contained: 15 uL KAPA HiFi Hotstart ReadyMix, 6 uL of combined i5 and i7 indexes (3 uL of each index, if they are being added individually), and 9 uL of PCR1.

Reaction conditions for the indexing PCR consisted of:

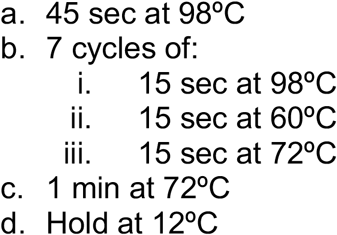

We then pooled equal volumes (5 uL) of each sample into a pool (to a total volume of 40 uL). We then cleaned the reaction using 1.7X SeraPure SPRI-beads (i.e. 68uL of bead solution), washing with ethanol, and eluting in a final volume of 40 uL of water. We checked the pool by TapeStation (D5000; Figure 1) and qPCR and sequenced it on a MiSeq NANO (500 cycle, i.e. 250 × 250 bp reads) at the Penn State Huck Genomics Core facility.

**Figure 1.**
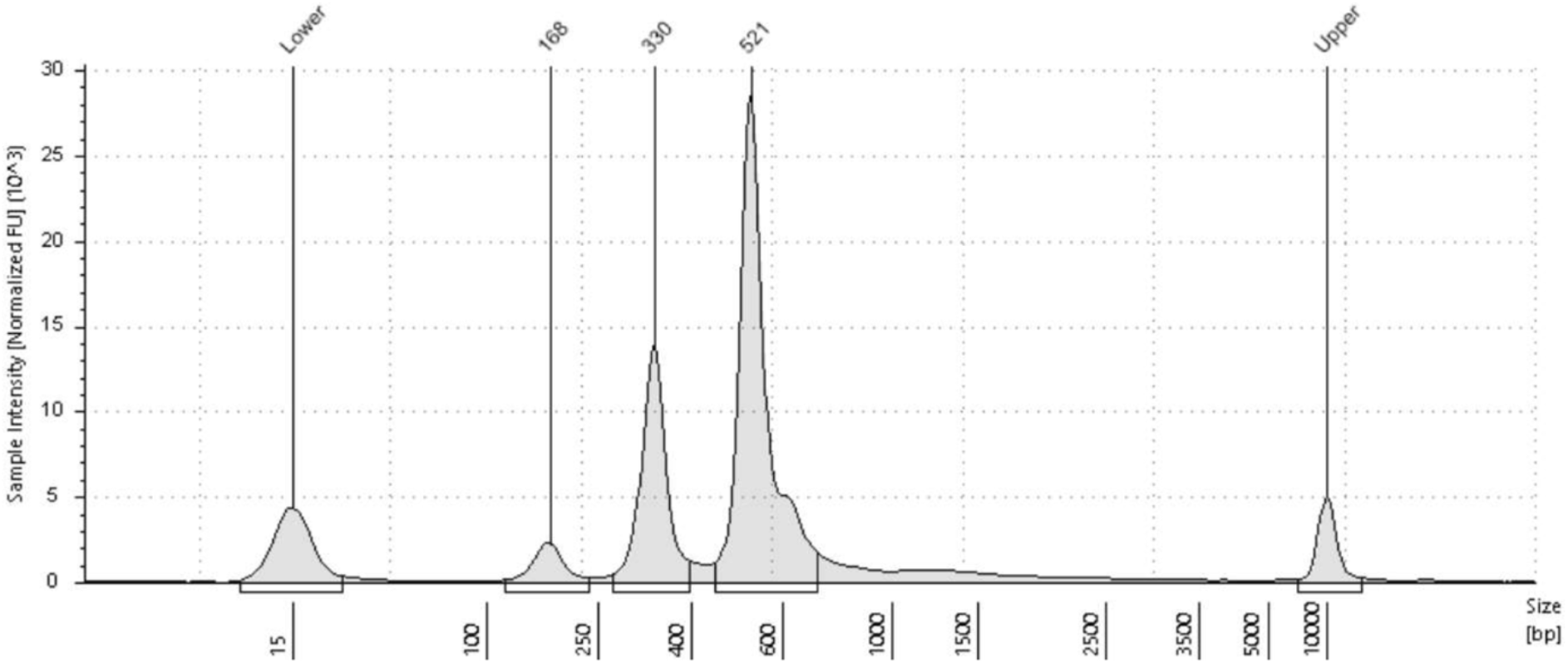
TapeStation results from pooled amplicon sequencing of 8 *Vermivora* warblers. The 521bp peak represents the main 4 amplicons, the small peak to the right is the 580 amplicon, and the smaller peak at ∼330 is the 299bp amplicon. The 168 peak is likely adapter dimer.

**Figure 2.**
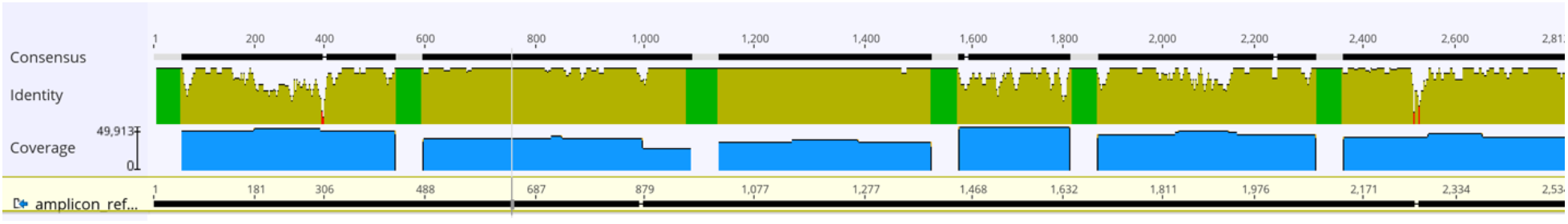
Coverage output from Geneious of sample BW 23-116. The green areas of the “identity” plot represent the ambiguous nucleotides seperating the 6 amplicon sequences.

### Bioinformatics

We created a single scaffold as a reference that consisted of six contigs, covering the six plumage loci, each separated by 50 ambiguous nucleotides “N’s” to aid alignment. We aligned the resulting reads to this scaffold using *Bowtie2* with the “--very-sensitive-local” set of presets. We then used *Samtools* to convert .SAM to .BAM files, sort, and index each alignment.

We called SNPs on each individual alignment using GATK’s (v 4.6.1.0) HaplotypeCaller, with the ERC flag set to “GVCF” output and output mode to “EMIT_ALL_CONFIDENT_SITES”. We forced GATK to call SNPs at only our focal SNPs using the -L flag and supplying it with the locations of the SNPs in a .bed file (available at https://github.com/david-toews/vermivora_amplicon).

Typically we would use GATK to called genotypes jointly across all “g.vcf” files, however even at the stage of combining genotypes GATK had issues with inserting missing data when the raw genotypes had high coverage. The extremely high coverage of the amplicon data may have produced issues at this stage, thus we simply extracted the raw genotypes from the individual .vcf files using a custom script (available at the GitHub repository).

## Results and Discussion

The pooled library resulted in three peaks on the TapeStation, one of which (at 168bp) was likely adapter dimer (Figure 1). The peaks above 300bp represented a size distribution that matched the predicted sizes of the six amplicons.

Sequencing resulted in approximately 140,000 reads per sample, which is much higher than is be necessary for calling genotypes. A target coverage is likely be between 10-50,000 reads per individual for future applications based on previous experience with meta-barcoding datasets, though even lower may be possible.

Genotypes across all six plumage associated loci match expectations based on RFLP cut sites and known phenotypes. In particular, the locus on Chromosome 20 is perfectly associated with the black throat color phenotype observed in all four golden-wings in this small sample (where a single heterozygous genotype would result in a plain, white throat). This locus includes the *ASIP* gene involved in the melanogenesis pathway (Toews et al. 2016, Baiz et al. 2020). Heterozygous bases were also seemingly correctly called by the pipeline—based on manual inspection of the raw alignments— such as the heterozygous site (Table 1) also clearly identified in Figure 3 in sample GW-23-155.

**Table 1.**
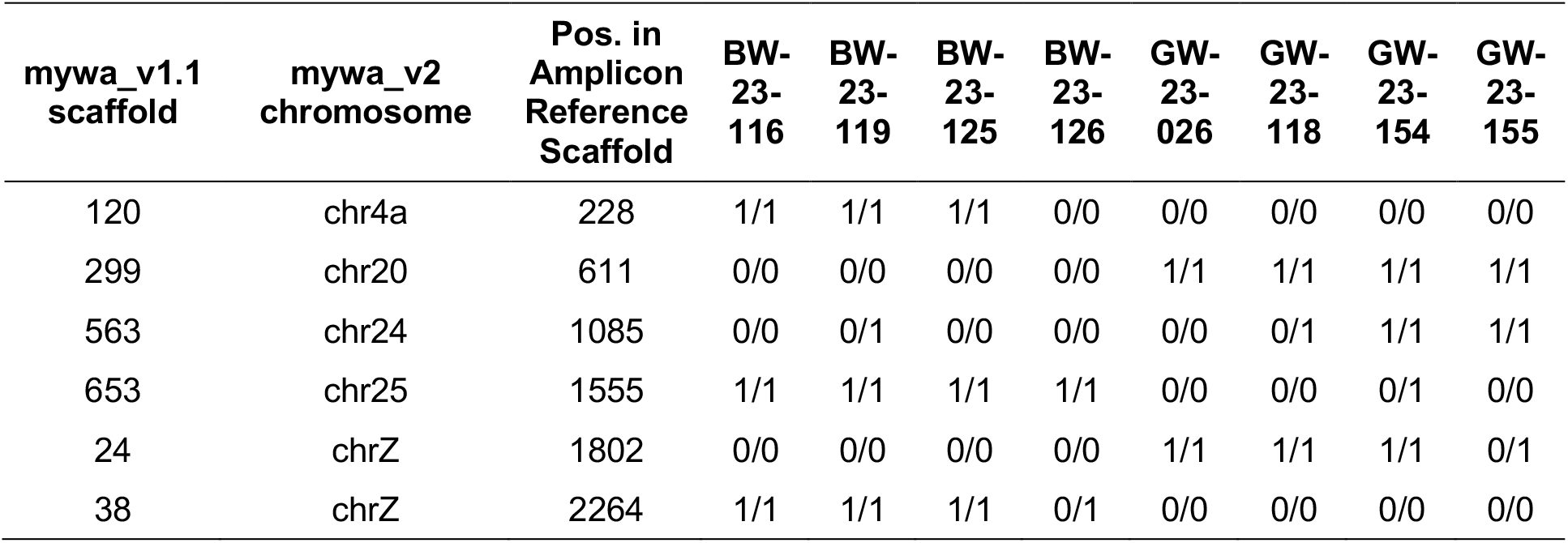
Genotype results from amplicon sequencing approach across four blue-winged warblers (“BW”) and four golden-winged warblers (“GW”). The sites are relative to three references: the scaffolds described in the original Toews et al. (2016) paper, the “v2” reference from Baiz et al. (2021), and the position in the artificial reference scaffold generated for this analysis.

**Figure 3.**
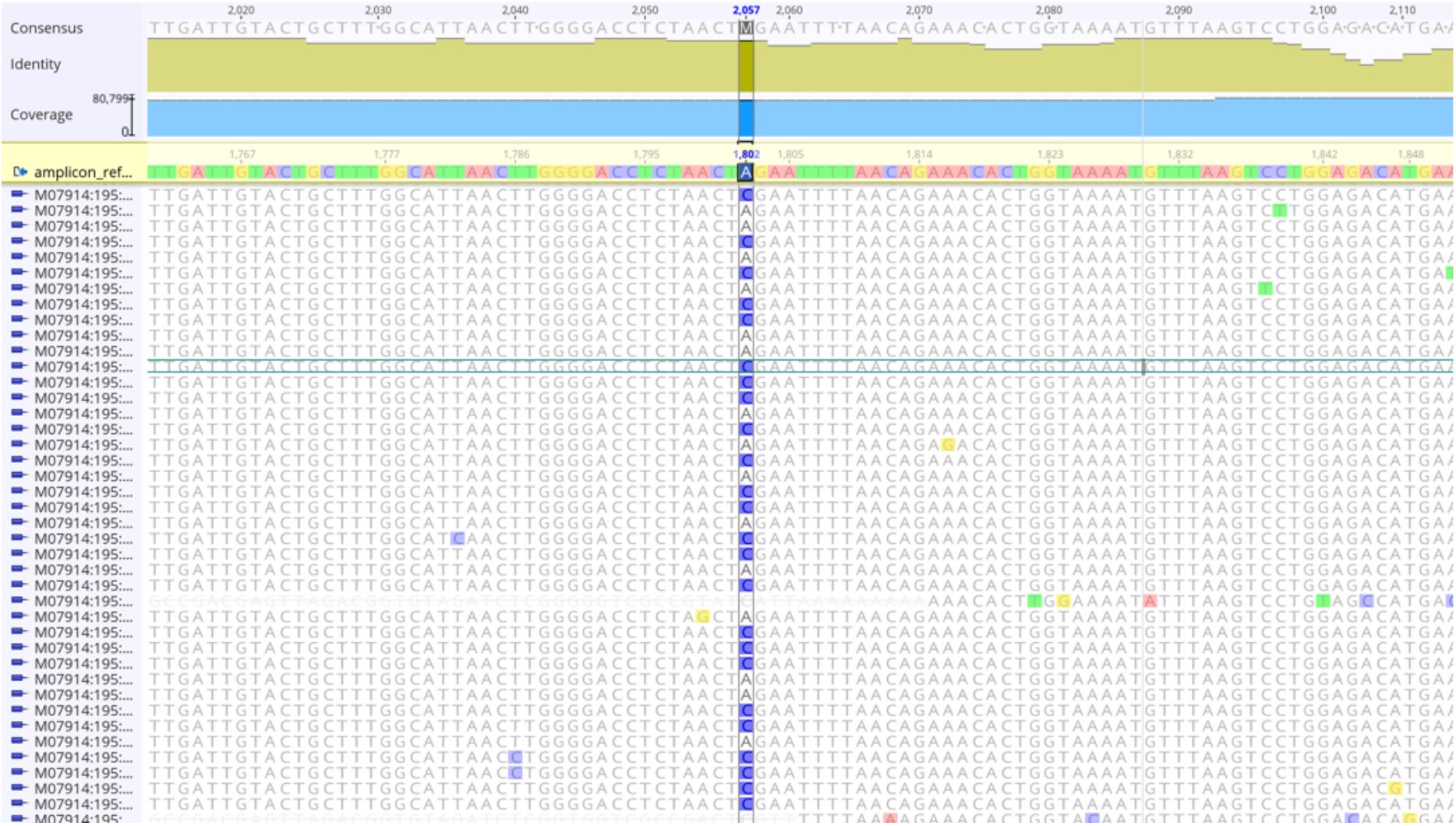
A heterozygous at site 1,802 bp for sample GW 23-155.

Multiplexing many more individuals is feasible and we have plans to include 384 individuals in a single run using a NextSeq P1 chemistry (600 cycles i.e. 300 × 300 bp), likely resulting in >250,000 reads per individual, as well as adding additional loci. Therefore, this appears as an efficient and cost effective method for unambiguously genotyping *Vermivora* at an important set of SNP loci across a large number of individuals simultaneously.

## Acknowledgements

Kurt Ongman and Amber Roth provided the samples used in this preliminary run. Methods were developed with inspiration and/or discussions with: Leonardo Campagna, Jennifer Walsh, Bronwyn Butcher, and Simon Kraemer. Computations for this research were performed on the Pennsylvania State University’s Institute for Computational and Data Sciences’ Roar Collab supercomputer. Research was conducted under Penn State’s PROTO201900879. Funding was supported by NSF DEB-2337828 and Pennsylvania State University, including funds from the Eberly College of Science and the Huck Institutes of the Life Sciences.

